# Genomic insights into adaptation to eco-regional and cultural variables across human populations from North, Central and Southeast Asia

**DOI:** 10.64898/2026.01.12.699033

**Authors:** Johanne Adam Doucet, Romain Laurent, Chan Leakhena Phoeung, Choduraa Dorzhu, Tatyana Hegay, Raphaëlle Chaix, Evelyne Heyer, Laure Ségurel

## Abstract

Natural selection has been extensively studied in humans, providing many examples of how climate, diet, and pathogens can translate into local selective pressures and thus phenotypic diversity across populations. However, most studies have focused either on a global scale, with a limited number of populations per geographic area, or on a very local scale. To complement these approaches, we collected and studied a large genomic dataset at a continental scale covering North, Central, and mainland Southeast Asia, and consisting of 863 individuals from 25 culturally diverse populations. We aimed to decipher the selective pressures across the Asian continent, comparing populations from different geographic areas and having contrasted subsistence strategies, using both intra-populations (iHS) and inter-populations (Fst) statistics. We detected both local and continental-wide signals, with some geographically and culturally complex patterns emerging when being compared to the literature. Interestingly, among the regions detected as significant only in mainland Southeast Asia, we found two of them clearly pointing to selective pressures associated with immunity. The first one, detected with iHS only, contains three peptidoglycan binding genes (*REG1A*, *REG1B* and *REG3A*) interacting with bacterial cell walls. The second region, the most significant one when intersecting iHS and Fst, contains *PELI1*, previously shown to modulate the immune and inflammatory response and to be under selection in Han Chinese and Oceanians. In addition, in North and Central Asia, we uncovered a region overlapping with *PTPRC*, a gene associated with viruses recognition, as well as another region including *GPHB5*, associated with lipid and carbohydrate metabolism. Interestingly, we further identified a signal on *PTPRG*, where variants have previously been associated with alcohol flushing in East Asia, not only in farmers but also in herders and hunter-gatherers, challenging the hypothesis that this phenotype was selected for in response to the transition to agriculture. In turn, when focusing on signals specific to given subsistence strategies, we found two genes related to immune functions (*TUBA1B* and *HERC1* ) exclusively in herders. In conclusion, while many signals of selection likely remain to be uncovered in less studied human groups, deciphering whether they are linked to immune, dietary or climatic factors is challenging, as subsistence strategies often covary with climate, which further influence pathogenic loads.

## Introduction

Over the last 50,000 years, humans have colonized a vast range of ecosystems, encountering diverse climates and interacting with different sets of pathogens. Furthermore, they have diversified their lifestyles, especially in the last 10,000 years, notably leading to contrasted dietary regimes. These changes resulted in genetic adaptations that have left distinct signatures in contemporary human genomes [1, 2]. Overall, eco-climatic factors, pathogenic load and dietary shifts have been repeatedly identified as important selective pressures driving a number of adaptations in humans [3, 4, 5, 6, 7, 8].

Focusing on the Asian continent, adaptation to cold environments has for example been found in Siberian populations in *UCP1*, which enhances non-shivering thermogenesis for heat production [9, 10, 11]. Adaptations to regional variation in ultraviolet (UV) radiation has also been studied in East and South Asians, with several genes involved in skin pigmentation identified as targets of positive selection, such as *OCAS2* and *DCT* central to melanin biosynthesis. *OCAS2* variants are associated with lighter pigmentation [12, 13], while mutations in *DCT* influence the shade and tone of pigmentation. *SLC45A5* and *SLC24A5* were also identified as targets of positive selection, with mutations modulating the efficiency and quantities of melanin produced, respectively [14]. The impact of extreme eco-climatic environments such as the Himalayan high plateau has also been characterized in Tibetan and Nepalese populations, with genes such as *EPAS1* and *EGLN1* contributing to more efficient oxygen assimilation in hypoxic environments by limiting excessive erythropoiesis [7, 15, 16, 17, 18] and modulating the degradation of hypoxia-inducible factors [19, 20, 18], respectively.

Pathogenic pressures, on the other hand, have been shown to result in positive selection of numerous variants in malaria-endemic regions such as Southeast Asia, South Asia and West Africa [21]. A study contrasting northeastern (from Japan, Korean and China) and southeastern (from the Philippines, Malaysia, Indonesia, Thailand, Singapore) Asian populations further highlighted local adaptation signatures at immunity-related loci, such as *DAPP1*, *IFNG* and *LEPR*, pointing to a geographically-structured adaptive response to immune pressures across Asia [22]. Adaptations to ancient viruses dating back around 20,000 years ago have also been identified in East Asians (Han Chinese, Japanese and Vietnamese) [23].

Finally, adaptations to diet have also been underlined, with genes regulating the metabolism and thus bioavailability of elements like selenium (*DIO2*, *GPX1* ) [24, 25, 26] and zinc (*SLC30A9*, *SLC39A8* ) [26, 27] being under positive selection in Han Chinese and Japanese populations. In Siberian populations (Yakut and Nganasan), mutations on genes linked to lipid metabolism (*PLA2G2A*, *PLIN1* and *ANGPTL8* ) show signs of positive selection [28]. These mutations likely optimize fat use and storage, as it is an important part of subArctic diets [29]. Other genes, such as *FADS1*, *FADS2*, *HADHA* and *HADHB* have also been shown to accommodate high-animal-fat diets and improve metabolic efficiency in fat-rich diets in Siberian and Inuit populations[30]. Additionally, variations in *CPT1A* optimizes dietary fat use for heat in Northeast Siberian populations (Chukchi and Koryat) [31].

In other cases, the nature of the selective pressure is unclear. For example, the 370A allele in *EDAR*, under positive selection in East Asian populations (Han Chinese, Japanese, Korean, Cambodian and Mongol populations) [32], is associated with traits like thicker hair and increased sweat gland density [33]. It has been hypothesized that this variant may have provided benefits in thermo regulation [32, 34], protection against pathogens [35] or improved vitamin D delivery in breastfeeding [34, 36], but the exact selective advantage remains unknown. The strong signature of positive selection on *ADH1B* (Arg47His, rs1229984) and *ALDH2* (Glu504Lys, rs671), implicated in ethanol oxidation and acetaldehyde breakdown, in Han Chinese, Japanese and Korean populations [37, 38, 39] is another example where the selective drivers are still debated. The mutations under selection lead to increased alcohol sensitivity and have been dated to appear around the same time as rice farming, supporting an hypothesis linking these adaptations with fermented rice beverages consumption [40, 41]. However, other hypotheses have been proposed, such as protection against excessive alcohol intake and its related diseases [42, 43, 44], leveraging the antimicrobial effects of acetaldehyde against infectious diseases or accessing clean drinking water [45, 46, 47].

Importantly, while we can see that numerous studies have previously aimed at identifying genomic signatures of natural selection in humans, there is a great discrepancy between populations, with most studies examining adaptive events at a large scale (across continents) by including only few East and South Asians, namely Han Chinese, Japanese, Vietnamese and Pakistani [48]. In parallel, multiple very localized studies have emerged, but do not include a comparison of their signals in a broader set of populations. Although Southeast Asia has recently gained more representation in public datasets [49], many areas, such as Central Asia, remain mostly unexplored. As a consequence, it is often challenging to correctly interpret the identified adaptive event and decipher whether they are driven by lifestyle, local climatic conditions, or the local set of pathogens.

To address this and better characterize the selective pressures associated with each adaptive signal, we collected and studied a large genomic dataset of 863 individuals across Asia, encompassing 25 populations with contrasted subsistence strategies and eco-climatic conditions. These populations mostly correspond to two regions, North and Central Asia (NCA) and mainland Southeast Asia (SEA). NCA is characterized by harsh continental climates, with extreme temperature fluctuations. The region is marked by large areas of steppe and desert, with relatively low rainfall and sparse vegetation, challenging thermo-regulation and metabolic efficiency. In addition, the northernmost parts of this region, extending into Siberia, are defined by subarctic environmental conditions, i.e. prolonged cold winters and short growing seasons, further intensifying pressures on energy balance, cold adaptation, and dietary resilience. In this region, pastoralism has been dominant, with many populations relying on meat and dairy products, providing high protein and fat intake [50, 51]. Some Central Asian groups however still retained their hunter-gatherers lifestyle until very recently. Some populations have also transitioned to agriculture during the Neolithic, and farming in this region mostly rely on crops such as wheat, barley, and millet [52, 50]. In parallel, SEA presents a tropical climate with high humidity, abundant rainfall, and monsoon seasons. Dense tropical forests and coastal ecosystems contribute to a different set of putative selective pressures, mostly related to pathogen resistance, heat tolerance, and humidity regulation. Most of the sampled SEA populations practice horticulture, with a diet mostly relying on dry rice [53]. In this study, we aimed at detecting global and local adaptive events in NCA and SEA populations, using both an intra-population (iHS) and an inter-population (Fst) approach. These statistics capture different aspects of positive selection: iHS identifies long haplotypes that are indicative of recent positive selection and does not require adaptation to be local; in contrast, Fst measures population differentiation at individual loci (and is thus less sensitive to low SNP density) and remains significant even if the selective sweep is over. To maximize the signal to noise ratio, we used a sliding windows approach, allowing us to focus on signals supported by multiple adjacent SNPs.

## Results

### Population sampling and genetic structure

We obtained genome-wide SNP data for 25 human populations from Asia: 12 populations from North and Central Asia (NCA, sampled in Uzbekistan, Kyrgyzstan and Russia) and 13 populations from mainland Southeast Asia (SEA, sampled in Laos and Cambodia), summing up to a total of 863 individuals. While most SEA populations correspond to horticulturists, we refer to them as farmers because their diet consist of high-starch diet. NCA populations present some variability in their ancestral subsistence strategy, with 3 farmers, 3 hunter-gatherers, and 6 herders. All 12 NCA populations were genotyped on an Illumina Omni1 array; 9 of the SEA populations were genotyped on the same array, while 4 of them were genotyped on an Illumina Omni2.5 array (Table S1). After filtering, we obtained an average of 607,753 and 957,999 SNPs per population, for each Illumina array, respectively.

The sampled populations form two distinct genetic clusters that clearly separate NCA populations from SEA on the first axis of the PCA (corresponding to 41% of the variation). The PC2 axis (7.4% of the variation) mostly separates eastern from more western populations within NCA (Figure 1). Furthermore, we observe an additional structure within NCA, with populations clustering based on their ancestral subsistence strategy. Indeed, in the eastern group, hunter-gatherers cluster separately from herders, and in the western group, farmers cluster separately from herders (Figure 1).

**Figure 1:**
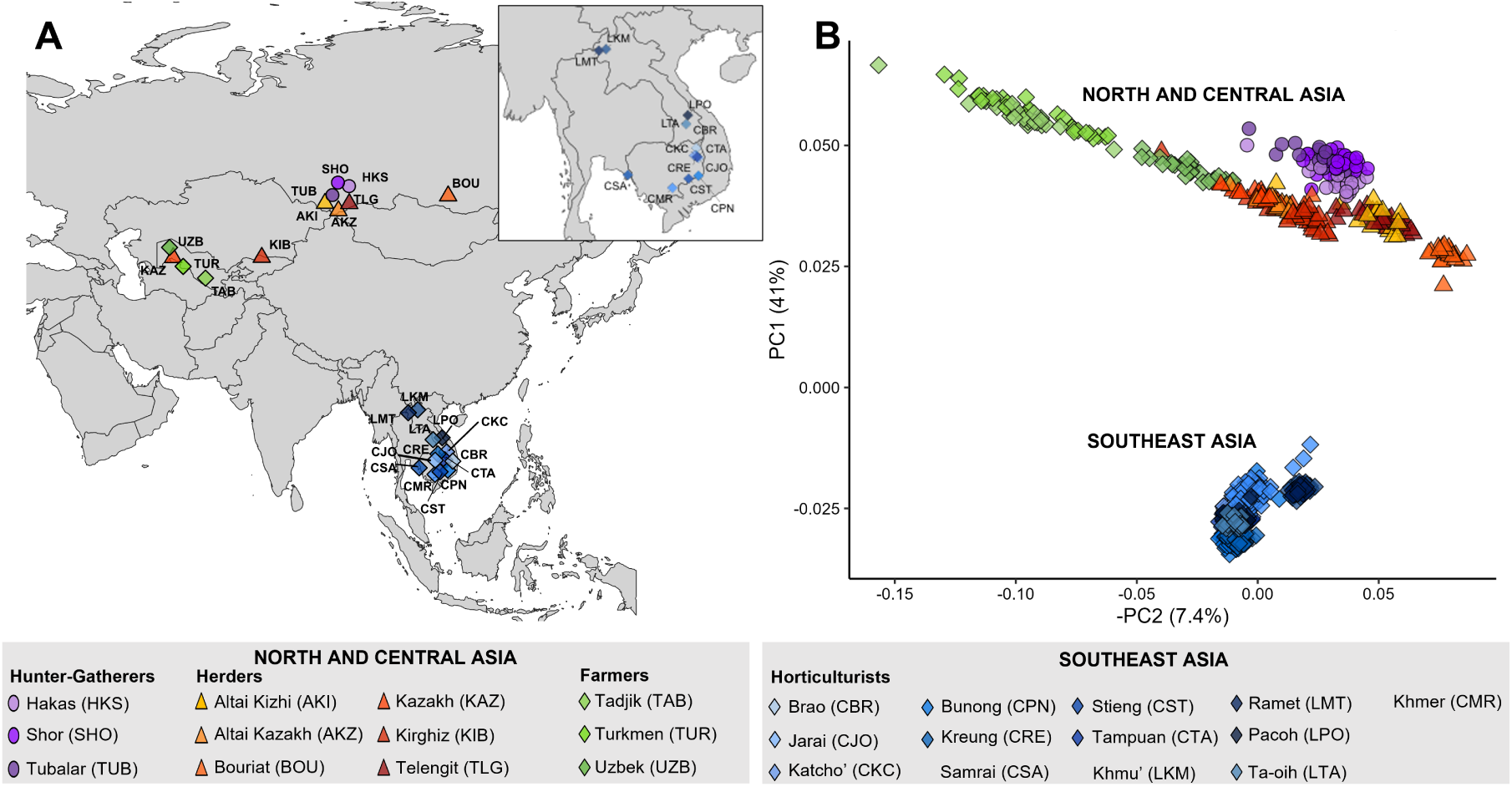
(A) Geographical map of the populations included in this study. **(B)**: Principal Component Analysis (PCA) on genome-wide SNP data, as performed using PLINK –pca [113].

### Signals of positive selection shared across populations

Our first objective was to detect which genomic regions show evidence of positive selection across Asian populations, so we calculated the iHS score [54] for each SNP in each population. We used a sliding window approach and defined significant SNPs as the ones having iHS scores within the top 1% of the genome-wide distribution, and significant windows as the ones having a proportion of significant SNPs within the top 1% of all windows. We identified in average 493 significant windows per population (min=475-max=513), with no significant differences between genotyping arrays (see Material and Methods, Figure S1). To focus on the most reliable signals, we retained only significant windows shared by at least two populations. We then concatenated the adjacent significant windows, resulting in a dataset of 1212 iHS-significant windows, with an average length of 158kb (Table S2).

To determine the most widespread adaptive events across Asia, we focused on the windows shared by most populations. We identified four genomic windows shared by at least 17 out of the 25 Asian populations (Table 1, Figure S2, Figure S3, Table S3). All of them were detected in approximately equal proportions in NCA and SEA (at least 7/12 NCA and 10/13 SEA populations). These windows range from 600kb to 1Mb and two of them are among the largest present in the overall dataset (Table S3, S4). A first windows includes *ERBB4* (Figure 2B), a second one includes both *EXOC6* and *CYP26A1* (Figure 2D), and a third one is close to *CNTNAP5* (Figure 2A). *ERBB4*, *EXOC6* and *CNTNAP5* have all previously been identified as some of the strongest selective signals in East Asia (including China, Japan, Cambodia and Russia) [55]. The fourth window, located on chromosome 2, encompasses multiple long non-coding RNA (Figure 2C), including *LINC0217* previously identified as a selection signal shared by multiple lowland but not highland Papuan populations [56].

**Figure 2:**
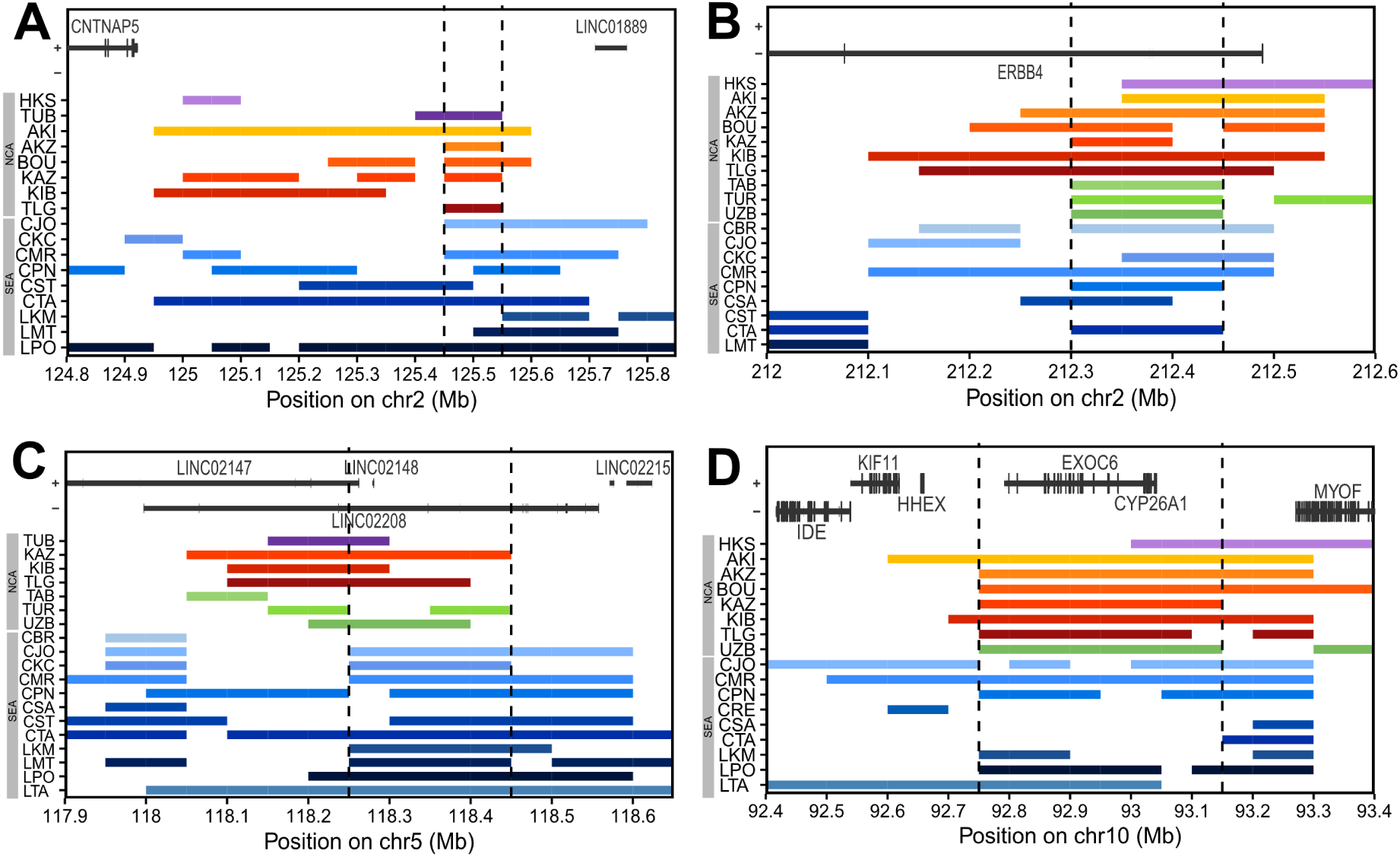
Illustration of the four continental-wide iHS signals, i.e., found in both NCA and SEA. Each colored rectangle represents the genomic window identified as significant in each population (which codes can be found in Figure 1). The positions of overlapping genes are represented on the top. The dashed lines represent the part of the region strictly shared by more than 11/25 populations. Only the populations contributing with at least one window are represented.

**Table 1:**
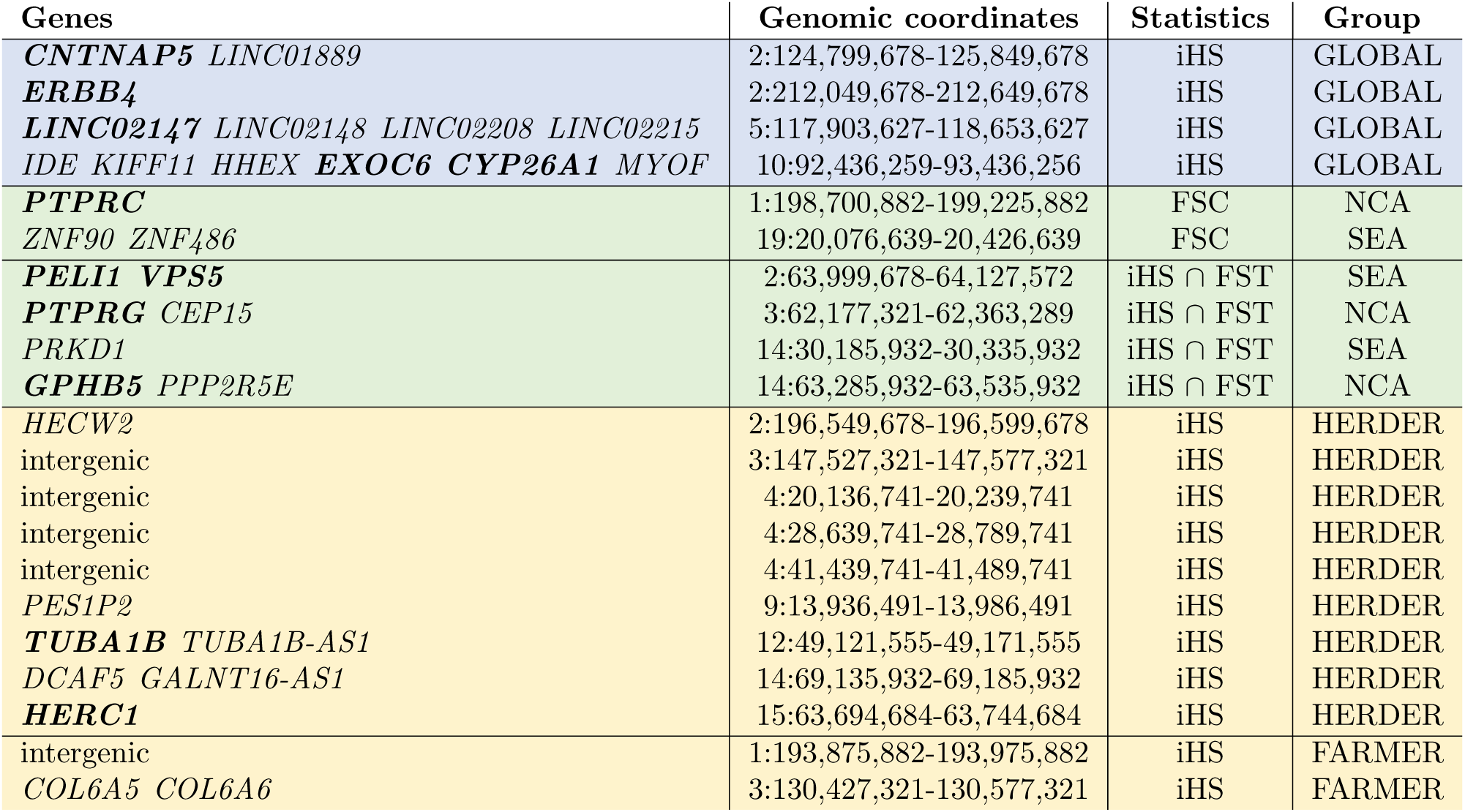
List of the main adaptive signals found in our study. The statistics used to detect these signals and the geographical or cultural scale at which they were identified is indicated. Genomic coordinates are in hg38. Genes in bold are the ones mentioned in the main text.

### Ecoregional adaptations: contrasting North and Central to Southeast Asians

We then asked how selection differs between NCA and SEA populations, to detect ecoregion-specific signals. We first performed Mann-Whitney rank tests comparing, for each of the 1212 iHS-significant window, the proportion of iHS-significant SNPs between NCA and SEA (see Material and Methods). After correction for multiple testing (FDR), we detected a total of 71 significant windows: 41 with a higher proportion of significant SNPs in NCA and 30 with a higher proportion in SEA (Table S5). These windows have an average length of 224kb, so tend to be slightly longer than the whole set. We then performed a functional enrichment test for NCA-specific and SEA-specific significant windows. No molecular pathway was found to be overrepresented in NCA. In SEA, peptidoglycan binding genes were found to be significantly overrepresented, due to a 450kb region containing three of them (*REG1A*, *REG1B* and *REG3A*), when 0.01 were expected (q-val= 1.2e-5) (Figure S4).

We next examined which SNPs most clearly separate NCA from SEA using Weir and Cockerham’s Fst. To this end, we merged the data from the Omni1 and Omni2.5 arrays, resulting in 329,203 SNPs. For each SNP, we averaged all Fst values contrasting one NCA to one SEA population (Figure S5). Similarly to the iHS approach, we determined Fst-significant SNPs and windows using a 1% threshold. Doing so, we identified 144 significantly differentiated windows between NCA and SEA (median length = 250 kb) (Table S6). We did not find any significant functional enrichment among the genes located in these windows. One of the top 10 Fst-significant windows encompasses three genes (*TRAFD1*, *NAA25* and *HECTD4* ) and has previously been described as under selection in East Asians (Han Chinese, Japanese, Korean) [57, 58, 59].

Next, we asked which SNPs show evidence of both a recent sweep and a strong population differentiation, combining iHS and Fst into a single Fisher Score (Fsc) (see Material and Methods). Among the 1120 top 1% Fsc-significant windows, we focused on the two regions that were detected in 6 populations or more from one ecoregion, but were absent from the other one (Figure 3, Table S7). One of these regions encompasses *PTPRC*, a gene involved in the recognition of viruses such as HIV-1 [60], hepatitis C [61] and HSV-1 [62]. This locus is under selection in 7 NCA populations (2/3 HG, 2/6 Herders and 3/3 Farmers) (Figure 3E, Figure 3F). The second locus, under selection in 7 SEA populations, contains two ZNF coding genes that had not been previously identified as under selection in the literature (Table1).

**Figure 3:**
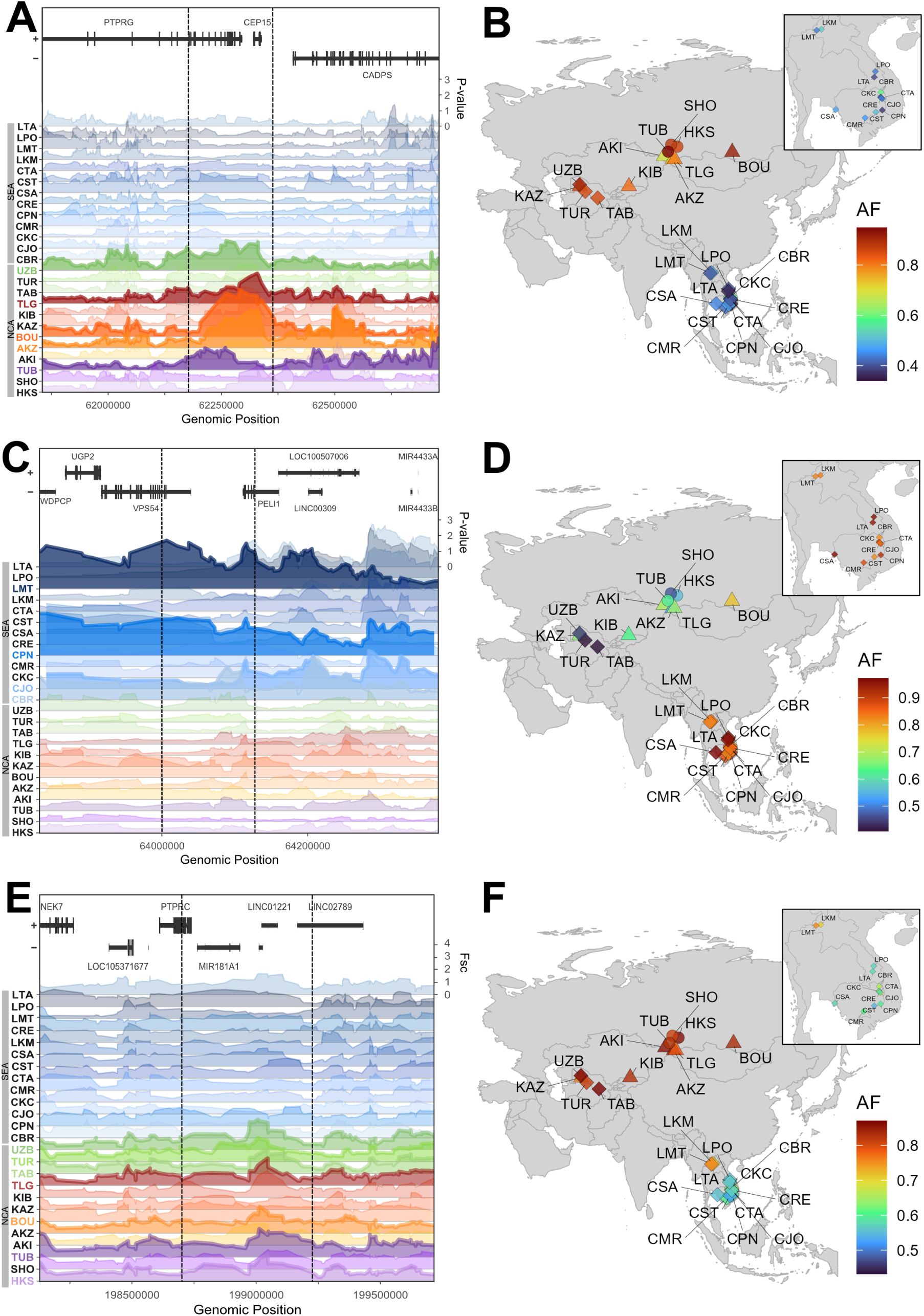
Illustration of three local adaptive signals contrasting North-Central and mainland Southeast Asia. **(A,C)**: The rolling mean (over 5 SNPs) of the pvalue-transformed iHS scores (ranging from 0 to 3.5) is shown for each of the 25 populations. **(E)**: The rolling mean (over 5 SNPs) of the raw Fsc scores (ranging from 0 to 4.5) is shown for each of the 25 populations. Panel A and E represents NCA-specific signals, and panel C a SEA-specific signal. Each row is overlapping with the neighboring rows. The dashed lines represent the significant region of interest. The population codes are colored when the window is found significant in the given population. **(B,D,F)**: Allelic frequencies in each of the 25 sampled populations for rs17066275 (*PTPRG*), rs329483 (*PELI1* ) and rs10800619 (*PTPRC* ). Each SNP was chosen for illustrative purposes and represents the one with the highest Fst score inside the interest region. For panel B, the frequency of the ancestral allele is represented, while for panels D and _1_F_5_, the frequency of the derived allele is represented, in accordance with the sign of the iHS score observed for each signal. Population codes can be found in Figure 1.

Given that the Fsc analysis can only be performed on the subset of SNPs common to all genotyping arrays, we sought an alternative approach to identify genomic regions showing both strong allelic frequency differentiation and within-population haplotype signals without losing too many SNPs. We thus looked at the overlap between the iHS-significant and the Fst-significant windows (71 and 144, respectively), and identified 4 such genomic regions (empirical pvalue=0.0001) (Table1, Figure 3, Table S8). Two of them correspond to a higher selective pressure in NCA, while the two others correspond to a higher selective pressure in SEA. One NCA-specific locus contains *PTPRG*, a gene linked to alcohol flushing in East Asian populations [39] (Figure 3A, Figure 3B). This signal is significant in 5/12 NCA populations, including 3/6 herders, 1/3 farmers and 1/3 HG populations. The other NCA-specific window contains *GPHB5*, a gene implicated in lipid and carbohydrate metabolism [63], present in 2/3 farmers and 1/3 HG (Table 1). As for South-specific signals, only one was found in a coding region. This window contains *VSP5* and *PELI1* (Figure 3C, Figure 3D) and was found significant in 4/12 SEA populations. Interestingly, *PELI1* has been identified in Han Chinese as one of the strongest immunity genes under selection [64] and also showed a moderately strong selective signature in Oceanians [55].

### Dietary adaptations: contrasting populations with different subsistence strategies in North and Central Asia

To uncover potential diet-related genetic adaptations, we contrasted populations with distinct subsistence strategies living in the same ecoregion, i.e. the 6 herders, 3 farmers and 3 hunter-gatherers from NCA. We performed Mann-Whitney rank tests comparing the proportion of iHS-significant SNPs between herders, farmers and hunter-gatherers. For each subsistence strategy, we contrasted populations from that subsistence strategy to all other populations (see Material and Methods). None of the genomic regions tested showed a significant effect after correction for multiple testing (FDR). To gain more power, we further examined which iHS-significant windows were consistently present only in herders, farmers, or hunter-gatherers, respectively. We found 3 genomic windows shared only by 3/3 farmer populations (empirical p-value=0,0004) (Table S9, Table S10), 9 windows exclusive to 4/6 herders (empirical p-value=0,0006) (Figure 4, Table S9, Table S10) and none shared exclusively by hunter-gatherers. No significant functional enrichment was observed for genes specific to herders or farmers. For herder-specific adaptations, four windows are intergenic, one contains a pseudogene (*PES1P2* ), and four contain genes, of which three had previously been shown to be under selection. Indeed, one window includes *DCAF5* and the antisense transcript *GALNT16-AS1*. While *DCAF5* has previously shown signatures of selection in the Lugbara (Nilo-Saharan farmers) [65], *GALNT16*, the sense gene corresponding to *GALNT16-AS1*, has been reported under selection in tropical forest populations [66]. The two other regions contain *TUBA1B* and *HERC1*, two genes previously linked to innate immunity [67, 68, 35].

**Figure 4:**
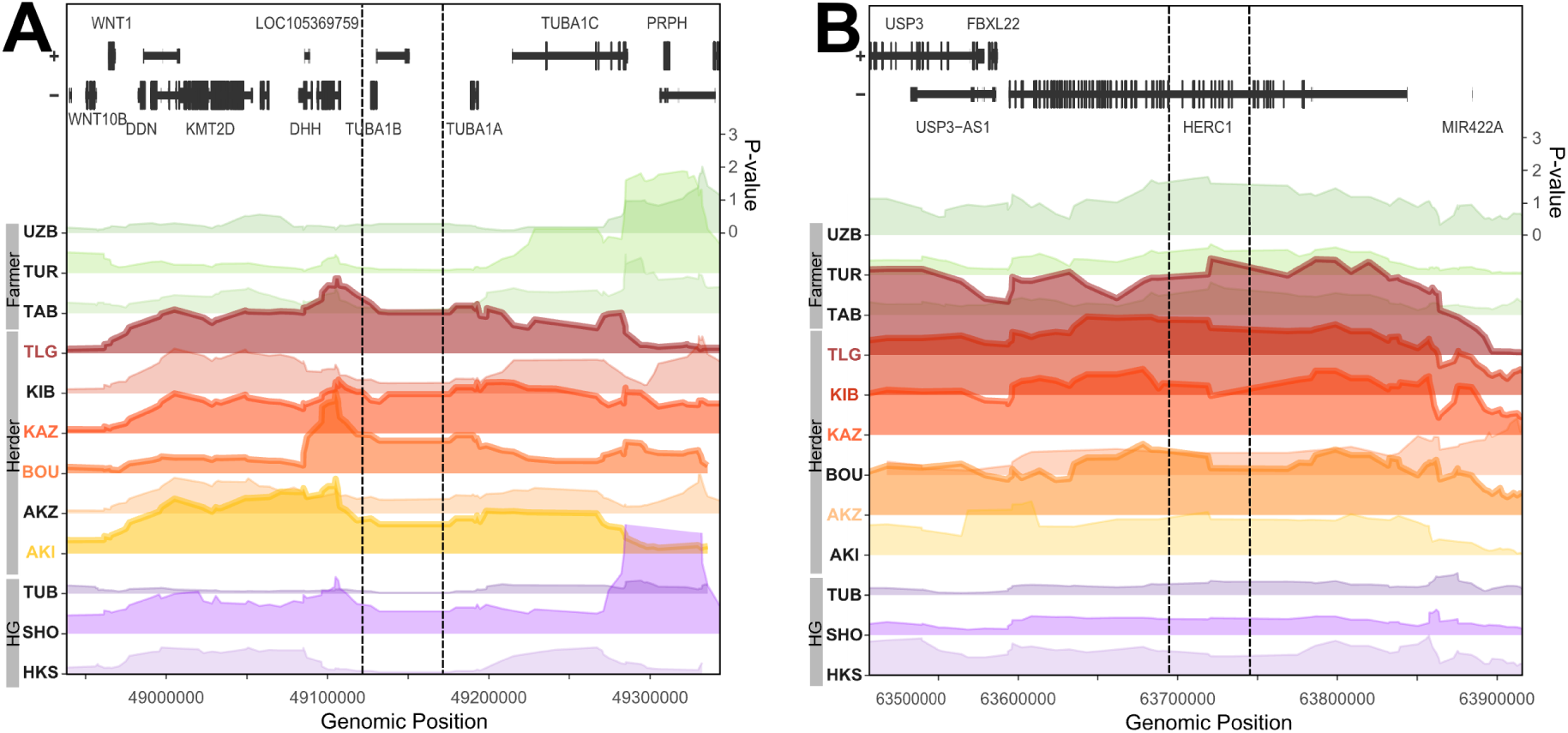
Illustration of two local adaptive signals associated with subsistence strategies in NCA. The rolling mean (over 5 SNPs) of the pvalue-transformed iHS scores is shown for each of the 12 NCA populations. Each row ranges from 0 to 3.5 and is overlapping with the neighboring rows. The dashed lines represent the significant region of interest. The population codes, which can be found in Figure 1, are colored when the window is found significant in the given population. Both panels are showing herder-specific adaptations.

## Discussion

### Methodological considerations

We combined iHS and Fst statistics to identify genomic windows where haplotype structure and allele frequency patterns both provide evidence for positive selection. To do so, we used two different methods: i) using a Fisher’ score (Fsc) and ii) intersecting the iHS and Fst outlier windows. Even though these methods use the same metrics, they combine them in different ways. Indeed, Fsc combines both statistics at the SNP level and thus restrict the analysis to SNPs shared by both genotyping arrays in our study. In practice, a SNP with a very high iHS score and an average Fst score (or vice versa) can result in a significant Fsc score. As a result, some significant windows are identified primarily due to one metric rather than a combination of both. In turn, our second method relies on identifying Fst-significant and iHS-significant windows separately, and then overlapping them. This method ensures that both statistics are significant for each window. However, both signals need not to be on the same SNPs, so there is a possibility that these windows encompass two different selective events that are physically close in the genome. These considerations might explain why the two windows identified with the Fsc are not among the top windows found when intersecting iHS and Fst. Therefore, it seems that these approaches are complementary to one another.

### Continental-scale adaptations across Asian populations

We first used iHS to identify the adaptive genomic regions that were shared by many populations at the continental scale and identified four windows shared by 17 or more populations (out of 25). The fact that three of these windows (containing *ERBB4*, *EXOC6-CYP26A1* and *CNTNAP5* ) had previously been identified in East Asians (China, Japan, Cambodia and Russia) in selective scans comforts the idea that they represent geographically broad (potentially Asian-wide) signals. The first window includes *ERBB4*, a gene known to be involved in the development of neural tissues, as well as heart and mammary gland development. It appears that the *NRG*-*ERBB4* pathway is under positive selection in all non-African populations, even though more strongly so in East Asians [55]. As of now, no adaptive phenotypes have been proposed to explain this strong selective event, calling for more experimental work on that gene. Interestingly, *ERBB4* and *EDAR* are both part of signaling pathways during the development of the mammary gland, even though they do not directly seem to interact [69, 70]. The second window contains *EXOC6*, identified as under selection in populations living in high-altitude (Tibetans and Nepalese) [71]. Functionally, the *EXOC6* protein plays a role in exocytosis, but no evolutionary hypotheses have been proposed so far to explain how *EXOC6* would contribute to high altitude adaptation. The same genomic window also contains *CYP26A1*, a gene classified as an ADME gene (genes related to Absorption, Distribution, Metabolism and Excretion) and involved in retinoic acid metabolim. It has been identified as highly differentiated between human groups [72]. *CYP26A1* and *EXOC6* along with *KIFF11* and *HHEX*, have been previously identified in selection scans in East Asian populations [55]. Overall, it seems that the geographical distribution of this adaptation is wider than initially found and spreads quite largely in Asia, which questions whether *EXOC6* is truly linked to high altitude adaptation and whether it is the target of selection in that genomic region. The third genomic region, neighboring *CNTNAP5*, has also been identified among the strongest XP-EHH signals in East Asians, together with a moderate selection in Papuans populations (Sepik and Puapan Highlanders and lowlanders) [55, 73, 74].

Our study revealed that this signal is also present in SEA populations and unexpectedly present in 8 NCA populations. The fact that these signals had previously not been identified in more populations is in part due to the incompleteness of public datasets such as 1KGP or HGDP that mostly focused on East and South Asia (China, Japan, Vietnam and Pakistan).

### Southeast Asia: pathogen-driven adaptations

Moving from continental to regional patterns, we focused on windows showing a spatially restricted distribution of selective events, i.e. being present specifically in SEA or NCA. Focusing first on SEA-specific adaptive events, we found various signals linked to immunity. Indeed, we identified an enrichment of peptidoglycan binding genes in this region, due to a window containing *REG1A*, *REG1B* and *REG3A* detected as significant in 8/13 SEA populations. Peptidoglycans are often present in bacterial envelopes and are a common target for innate immune cells [75, 76]. Secondly, the only SEA-specific adaptation we found in a coding region when intersecting iHS and Fst windows, containing *VSP5* and *PELI1*, is also related to immunity. The latter gene has indeed been identified in Han Chinese (CHB) as one of the strongest target of selection among immunity genes [64] and also shows a moderately strong selective signature in Oceanians (PNG) [55]. This comforts the view that SEA is a region with very high human pathogenic load [77]. Indeed, eco-climatic factors such as high humidity, high temperatures, monsoon seasons and overall broad zoonotic interfaces constitute pathogenic reservoirs. Interestingly, the adaptations we found in SEA seem to specifically be linked with bacterial pathogens, which are the most predominant type of infections in SEA in current times when excluding malaria cases [78]. Altogether, studying diverse populations in Cambodia and Laos allowed us to underline the need to further investigate immune adaptations unrelated to malaria, such as those against bacterial pathogens, in accordance with findings in [49].

Regarding malaria, it might be surprising that we did not find any evidence for selection on such genes in our analyses. Indeed, studies of ancient *Plasmodium* lineages has pointed to SEA as a long-term ecological niche for malaria [79], and multiple genetic adaptations have been found in SEA [80, 81] and other malaria-endemic regions [82, 83, 84], underlining the strength of this selective pressure [85]. This could be explained by the fact that variants associated with malaria resistance are often under balancing selection [86, 87, 88], which cannot be detected by our approach and/or because most of SEA populations sampled for this study live in the Cambodian highlands and Laotian mountains, a less favorable mosquito breeding environment [89]. Better characterization of malaria exposure in these populations would be of interest.

### North and Central Asia: immune and metabolic adaptations

We then looked at NCA-specific signals. Interestingly, results from the Fsc analysis highlighted *PTPRC*, a gene involved in the recognition of viruses such as HIV-1 [60], hepatitis C [61] and HSV-1 [64]. This locus is under selection in 7 NCA populations, more specifically in 2/3 hunter-gatherers, 2/6 herders and 3/3 farmers, revealing it is shared by all subsistence strategies. Furthermore, this gene has been shown to be under positive selection in Tibetan population [90], as well as in Maniq populations, a hunter-gatherer population from Thaïland [91] and in Amazon hunter-gatherer populations (Ashaninkas, Matsiguenkas, Matses and Nahua) [92]. This pattern could reflect a common pathogenic pressure acting across northern and southern populations, independently of eco-climatic region. Alternatively, it may result from distinct viral pressures in different populations, with NCA populations and tropical hunter-gatherers being exposed to different pathogens or environmental conditions that nonetheless target the same immune gene.

In addition, we identified *PTPRG* (Protein Tyrosine Phosphatase Receptor Type G) as a NCA-specific locus. This gene has been previously identified as a top signal for alcohol flushing in East Asians (Koreans and Chinese) when adjusting GWAS models for *ADH1B* and *ALDH2* [39]. Another study also implicated it in alcohol dependence [93] in European-American and Africa-American subjects. Interestingly, in our dataset, the populations harboring the strongest signal (iHS) at this locus are three herder populations. Even though the causal SNP associated with alcohol flushing was not on our genotyping array (and we found no SNPs in LD with it), this result is surprising in regard to the current hypotheses linking alcohol flushing to rice domestication and consumption of rice-fermented products [40, 41]. Alternatively, *PTPRG* has been proposed to be involved in immune signaling and modulation of inflammatory responses. Selection at this locus could thus be the result of pathogen interactions, an hypothesis that has also been proposed for *ALDH2* and *ADH1B* [45, 46]. In parallel, we found that one of the 10 most differentiated genomic regions between NCA and SEA contains *HECTD4*, a gene associated with alcohol consumption in East Asians [58], as well as metabolic traits linked to Type 2 Diabetes [59, 94]. It is however found on a long LD block next to *ALDH2*, making it arduous to know which is the actual target of selection [95, 58]. Altogether, these findings indicate the presence of selective pressures on or next to multiple genes involved in alcohol flushing in NCA, not only in farmers, but also in herders and hunter-gatherers, raising new questions about the origins of adaptation on this phenotype. Relatedly, another study identified strong selection on polymorphisms related to alcohol dependence in herders and hunter-gatherers from Northwest Asia (Khantys, Selkups, Nenets, Shor, Nganasan, Evenks, Evens, Sakha and Kets) [96] further supporting that alcohol-related selective pressures have also operated in non-agricultural populations.

We further found adaptive events clearly involved in dietary adaptation. We indeed observed adaptive signatures in *GPHB5*, linked to lipid and carbohydrate metabolic processes. *GPHB5* was found under selection in two NCA populations (1/3 Farmer and 1/3 HG). This locus has previously been identified as under strong selection in Europe, Middle-East and South Asia [55]. However, it has never been identified in East Asia. A study found that *GPHB5* can be used as a biomarker for metabolic syndrome (the presence of multiple risk factors that usually include obesity, insulin resistance, impaired glucose tolerance, low HDL cholesterol and high blood pressure) [63] and interacts mainly with genes involved in glucose and lipid metabolism, as well as energy balance.

Finally, we investigated adaptations unique to populations sharing the same subsistence strategy in NCA using iHS. We identified 2/9 herder-specific windows that point to immune functions: *TUBA1B* plays a key role in the MHC class II antigen presentation pathway, whereas *HERC1* has been linked to MHC class I and host–pathogen interactions. Interestingly, *HERC1* has previously been identified in selection scans in Chinese and Japanese populations, making it not exclusive to herders [55, 68].

Overall, the boundaries between adaptations driven by lifestyle or by environmental variables are often blurry. Ecoclimatic parameters such as altitude, UV radiation, temperature, precipitations or humidity influence the types of dietary resources in each ecoregion. As a result, subsistence strategies often covary with climate. For example, high-altitude populations face ecological challenges such as hypoxia, low temperatures, and increased UV radiation [97], but also correspond to low-fiber, high-fat and high-protein diets [98]. Similarly, Arctic regions have limited crop potential, favoring herding; arid and desert areas also promote pastoralism. Cultural practices thus interact with ecological constraints to determine what can be gathered, grown or bred, which often makes it difficult to determine whether genetic adaptations are primarily driven by dietary composition or environmental factors. Taking this into consideration underlines the importance of multi-layered approaches combining genomic, ecological, and cultural data, enabling a more nuanced understanding of the drivers of human adaptations.

## Material and Methods

### Dataset

Blood or saliva samples were collected in North and Central Asia between 2001 and 2012, and in Southeast Asia in 2012 and 2013. Informed consent was given by all participants. Detailed number of individuals per population are given in Table S1. The details for collection, molecular methods and preprocessing of genomic data are described in [99] and [100] for NCA and SEA, respectively.

### Liftover and quality controls

Genotyping arrays genetic coordinates were lifted from GRCh37 to GRCh38 using UCSC liftOver [101]. The reference allele was defined based on the human reference genome available at https://www.ncbi.nlm.nih.gov/assembly/GCF_000001405.26/. We filtered out indels and singletons, as well as variants with no position, duplicate positions and monomorphic SNPs. We excluded individuals with a call rate lower than 90% and variants with a call rate lower than 5%. Lastly, we removed variants in Hardy Weinberg disequilibrium (p-value *<* 10*^−^*^5^). All these steps were performed using BCFTools [102] or VCFTools [103].

### Phasing

We reconstructed haplotypes using SHAPEIT [104] with the 1000 Genomes Phase 3 genomes as a reference panel [105]. We phased our data twice with the same LD-based genetic map [106], as well as once with a pedigree-based genetic map [107], to investigate the robustness of the phasing process, as well as the effect of the genetic map, on our downstream analyzes. We looked at the impact of phasing on the unstandardized iHS (uniHS) and iHS values for each population. Spearman’s rank correlations coefficient computed on uniHS and iHS values (*̄x_uniHH_* (*r*^2^) = 0.994968 and *̄x_iHS_*(*r*^2^) = 0.993072) indicate a very good consistency of the iHS statistics between the two replicated phasing processes using LD-based map. The choice of the genetic map seems to play a more important, but still minor role on uniHH and iHS values (*̄x_uniHH_* (*r*^2^) = 0.9801 and *̄x_iHS_*(*r*^2^) = 0.9693). These differences are likely introduced by the difference in SNP density between the genetic maps, which influences the resulting SNPs density of the dataset.

### Intrapopulational metrics (iHS)

#### Data preparation and genome-wide scans

Once phased, the reference allele was defined as the ancestral one, based on the information obtained in [108] using VCFTools [103] and a customized script (available on demand). The final datasets contained respectively 755,695, 741,310, and 1,211,146 SNPs as well as 325, 368 and 143 individuals.

Genome-wide scans were performed using Selscan [109] on each population independently with the *–keep-low-freq* flag. The number of individuals in each population served as number of bins for the normalization step in the associated software *norm*.

#### Outlier extraction

We considered as significant the SNPs with an absolute iHS value within the top 1% of the whole genome distribution. We then computed the the proportion of significant SNPs in sliding windows of 100kb (with a step of 50kb). We extracted the windows having the 1% highest proportions of significant SNPs after discarding windows containing 5 SNPs or less. A set of significant windows were obtained independently for each population. We kept windows shared by at least 2 populations using bedtools *multiinter* [110] and concatenated them using a customized script (available on demand). On average, each window contained 35 SNPs independently of the genotyping array. The proportion of significant SNPs in each window was recalculated after concatenation. We verified that the proportion of significant SNPs per 100kb window was not different between populations or genotyping arrays (Figure S1).

#### Tests for local adaptation

Ecoregion: For each window, we contrasted, between populations from NCA and SEA, the median proportions of iHS significant SNPs using a Mann-Whitney-Wilcoxon test. This test was applied to each of the 1212 previously identified windows shared by 2 or more populations. After correcting for multiple testing (FDR method), we extracted the windows with q-values = 0.05. Subsistence strategies: For each window, we contrasted, between populations of a given subsistence strategy (farmers, herders or hunter-gatherers) and all other populations, the median proportions of iHS significant SNPs using a Mann-Whitney-Wilcoxon test. This test was applied to each of the 297 previously identified windows shared by 2 or more populations from NCA. After correcting for multiple testing (FDR method), we extracted the windows with q-values = 0.05.

We further developed a second method to identify the windows only found in some or all populations belonging to the same subsistence strategy. For each group, we calculated the expected number of such identified windows by chance, by permuting the population labels 10,000 times. Using the resulting empirical pvalue (Table S10), we excluded all windows identified as hunter-gatherer-specific and kept windows shared by 4/6 or more herders and 3/3 farmers.

#### Inter-population tests (Fst)

For measuring the Fst, we merged the different genotyping arrays into one dataset to which we applied the same quality controls as described in Material and Methods. We obtained a merged dataset of 329,203 SNPs and 863 individuals. Fst were computed pairwise using Vcftools –weir-fst-pop [103], and the average value between pairs of populations from NCA versus SEA was calculated for each SNP. We considered as significant the SNPs with a Fst in the top 1% of the genome wide distribution and computed the proportion of significant SNPs in sliding 200kb windows (step of 100kb). Windows containing less than 5 SNPs were discarded. Allelic frequencies were computed for all populations from our dataset using vcftools –freq on the SNP with the highest Fst score across all populations.

### Combining iHS and Fst

#### Fishers’ score (Fsc)

The Fsc was computed as the sum of log10(percentile rank iHS) and log10(percentile rank Fst) for each population. Here, the Fst was computed, for each population, by contrasting it to all other populations from the other region. We calculated the proportion of significant SNPs for each 300kb sliding windows (step of 150kb) and considered as significant the genomic windows in the top 1% of this distribution. These windows were then concatenated to constitute a final dataset of 1120 genomic windows shared by at least two populations. Then, we focused on windows that were shared by NCA-only or SEA-only populations in a significant way as described earlier for subsistence strategies. To assess significance, we computed the number of windows shared by chance for all combinations of any 2 to 13 random populations and repeated this 10,000 times. This gave us an empirical pvalue (Table S10). Based on that, we found that windows shared by 5 or less populations were not statistically significant and thus, we retained windows being significant and shared by 6 or more populations from NCA or SEA, respectively.

#### Overlapping iHS and Fst outliers

We then intersected the genomic coordinates of the 144 Fst significant windows (see Interpopulational tests (Fst)) with the 71 windows identified using non parametric tests (see Intrapopulational tests (iHS), local adaptation tests). Empirical p-value was computed by intersecting the genomic coordinated of 71 randomized windows drawn from the initial pool of 1212 with the 144 Fst significant windows. This was done 10,000 times.

#### Gene annotation and functional enrichment

Gene annotations were performed on 41 NCA, 30 SEA specific windows and 144 Fst significant windows, as well as for 9 herder-specific and 2 farmer-specific windows. For each, the genomic coordinates of the windows were intersected with GRCh38 human assembly using bedtools [110]. Only gene names were retained and uploaded to Panther[111, 112] where we performed a statistical over representation test.

## Acknowledgments

First of all, we thank all study participants, without whom this research would not have been possible. We are grateful to all the collaborators who contributed to the generation, preparation, and contextualization of the data used in this study, and who were already recognized as authors in previous publications of these datasets. For the NCA dataset, we thank Philippe Mennecier and Boris Chichlo for their contribution to field data acquisition, as well as Nina Marchi for her contributions to data acquisition, processing and pipeline implementation. For the SEA dataset, we thank Yves Buisson, Frédéric Bourdier, Olivier Evard and Samuel Pavard for their contributions to field data acquisition. We are grateful to Sophie Lafosse for her involvement in both fieldwork and wet-lab sample preparation. We also acknowledge Étienne Patin and Charlotte Antoine for valuable scientific discussions.

