## Supplementary Figures for "Genomic insights into adaptation to eco-regional and cultural variables across human populations from North, Central and Southeast Asia"

Johanne Adam Doucet[1], Romain Laurent[1], Chan Leakhena Phoeung[2], Choduraa Dorzhu[3], Tatyana Hegay[4], Raphaëlle Chaix[1], Evelyne Heyer[1], and Laure Ségurel[5]

[1]Muséum national d'Histoire naturelle, UMR 7206 CNRS Eco-anthropologie, Université Paris Cité, Paris, France

[2]Rodolphe Mérieux laboratory, University of Health Sciences, Phnom Penh, Cambodia

[3]State University of Tuva Republic, Kyzyl, Russia

[4]Republican Scientific Center of Immunology, Ministry of Public Health, Tashkent, Uzbekistan

[5]Université Lyon 1, UMR CNRS 5558 Laboratoire de Biométrie et Biologie Évolutive, Villeurbanne, France

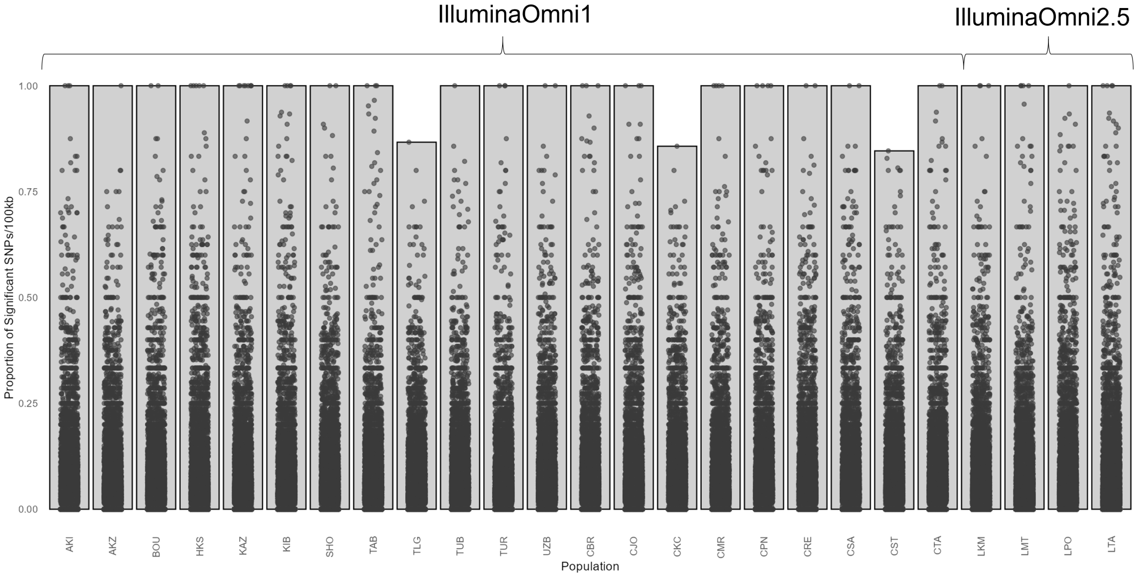


Figure S1: Proportion of SNPs with significant iHS in 100kb windows in the different populations. The genotyping array on which each population was genotyped is indicated at the top. The first 12 populations are from NCA, while the 13 last ones are from SEA.

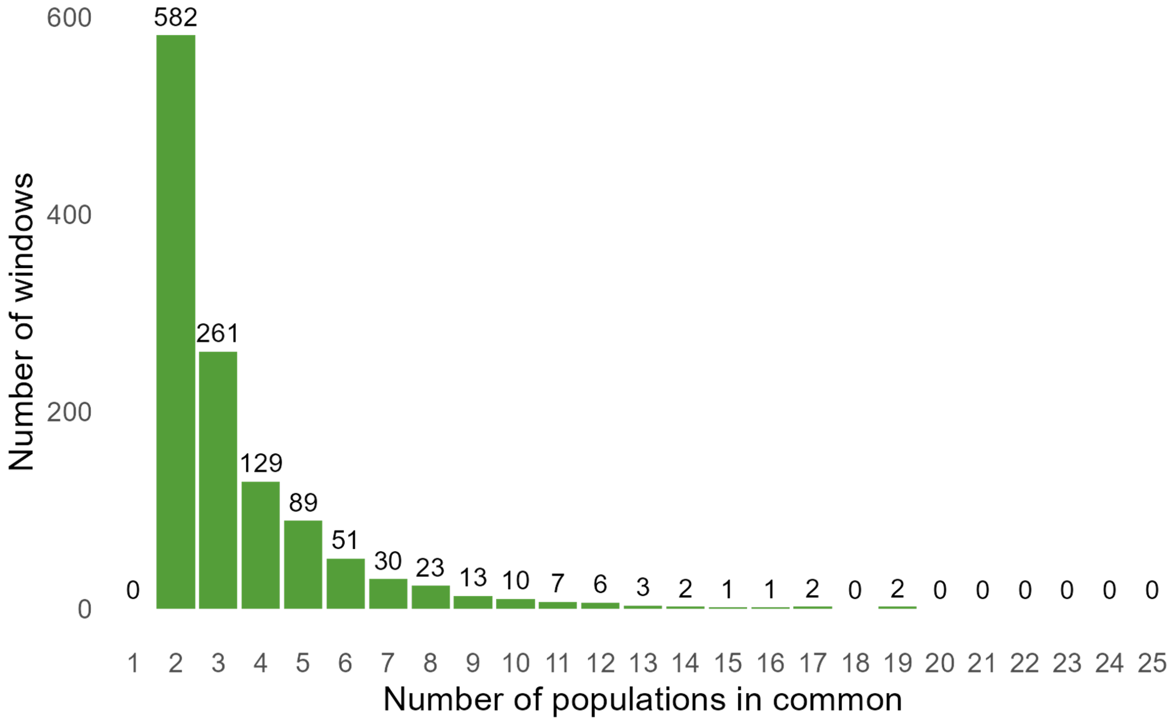


Figure S2: Distribution of the number of populations having iHS-significant windows (i.e., sharing a signal) across the 1212 windows detected as significant in 2 or more populations.

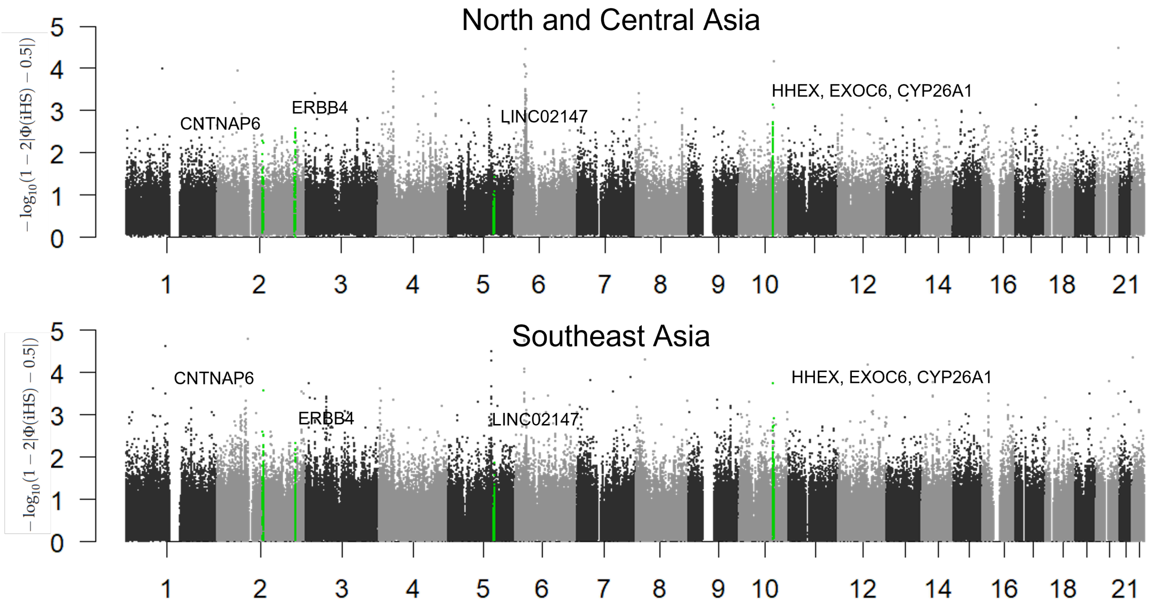


Figure S3: Manhattan plot of iHS pvalues in NCA (A) and SEA (B). Only the SNPs shared by all populations in either NCA or SEA were retained, and their pvalues were averaged for this illustration. Green regions represent the intervals shared by most described in Table 1 and in Figure 2.

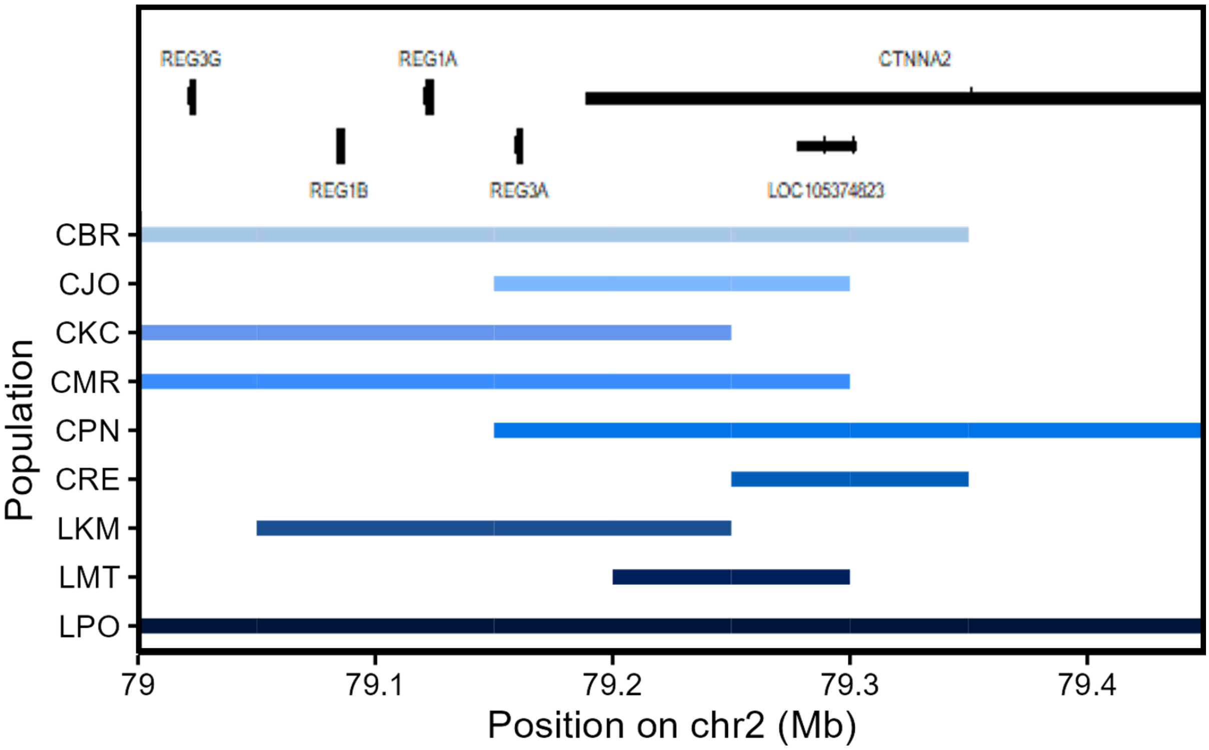


Figure S4: Illustration of the genomic window causing the functional enrichment in peptidoglycan-binding genes in SEA. Each colored rectangle represents the genomic window identified as significant in each population, without concatenation. The positions of overlapping genes are represented on the top. The population codes can be found in Figure 1.

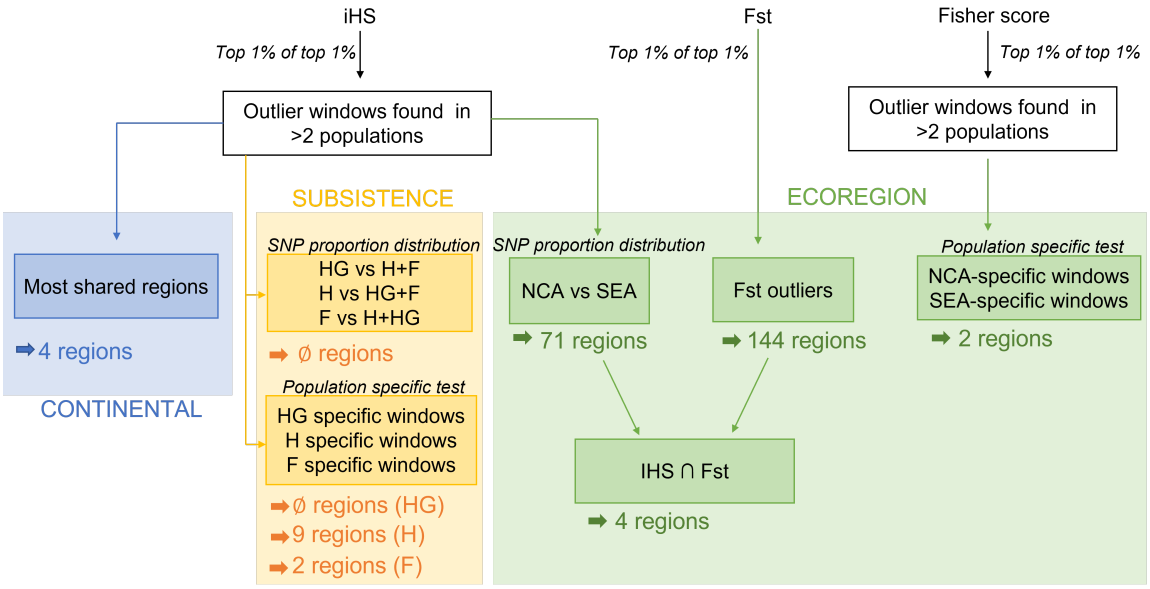


Figure S5: Overview of the methodological design and tests performed in this study, for each statistics used. iHS and Fsc were computed per population, whereas Fst scores were averaged over population pairs to obtain one metric differentiating NCA and SEA. The bold horizontal arrows indicate the number of genomic regions identified as under selection, for each layer of analysis. SNP proportion distribution refers to Mann-Whitney tests performed on the proportion of significant SNPs (iHS or Fsc). Population specific tests refer to windows being only found significant in populations belonging to the same group and not the others.
